# Single-disc optical visualization in photoreceptors uncovers protein architecture and compartmentalized pathology

**DOI:** 10.64898/2026.05.19.725875

**Authors:** Simone Mortal, Jesús Planagumà, Enric Pérez-Parets, Pablo Loza-Alvarez

## Abstract

Photoreceptor discs are densely stacked membranes housing visual pigments, yet their tight 35 nm spacing has kept molecular organization beyond the reach of optical microscopy. Here, we employ iterative ultrastructure expansion microscopy (iU-ExM) to achieve the first optical visualization of proteins within individual photoreceptor discs at 12 nm effective resolution through 20-fold expansion. We demonstrate that rhodopsin occupies 92% of the disc radial extent in situ, substantially exceeding the ∼50% area fraction reported from atomic force microscopy of isolated membranes. We provide the first optical detection of peripherin-2 within disc incisures and resolve connecting cilium and centriolar appendage architecture in three dimensions. Application to the RCS rat model of retinitis pigmentosa reveals compartmentalized pathology: outer segment disc spacing increased 29% while centriolar architecture remained preserved, suggesting intact protein trafficking during early degeneration. By bridging molecular identification with ultrastructural context, this work opens optical access to individual membrane compartments within densely packed cellular architectures previously resolved only by electron microscopy.

## Introduction

Vision in vertebrates depends on specialized sensory neurons called photoreceptors, which transform light into neural signals through phototransduction. This molecular cascade occurs within the outer segment, a highly modified primary cilium containing hundreds of tightly packed disc membranes that maximize photon capture. Each disc is a flattened membranous sac separated by approximately 30–35 nm, creating an expansive surface area optimized for visual sensitivity (*1–3*). The outer segment undergoes continuous renewal, with new discs forming at its base via evagination from the ciliary membrane while older discs are shed at the apex and phagocytosed by the retinal pigment epithelium (*4*).

Rhodopsin, the light-sensitive G protein-coupled receptor that serves as both photon detector and structural scaffold for disc membrane integrity, dominates the molecular architecture of photoreceptor discs. Early biochemical studies estimated that rhodopsin comprises approximately 90% of outer segment proteins, with mouse rod discs containing ∼75,000 rhodopsin molecules each (*5*). However, atomic force microscopy (AFM) studies on isolated disc membranes reported that rhodopsin forms paracrystalline nanodomains occupying only ∼50% of the membrane area (*6–8*). Clarifying this discrepancy is essential, as interpretation of disease-related changes in retinal degenerations would benefit from accurate baseline values for native rhodopsin organization in intact tissue.

Peripherin-2 (Prph2) represents another critical disc component, localized to the highly curved disc rims where it forms large oligomeric complexes with its homolog ROM1 (*3, 9, 10*). These tetraspanin proteins are essential for disc morphogenesis: peripherin-2 knockout abolishes outer segment formation, while its tetraspanin core induces membrane curvature required for disc rim formation and its C-terminus suppresses ciliary ectosome release (*10–12*). Beyond the external rim circumference, disc rims include incisures, longitudinal indentations thought to facilitate diffusion of signaling molecules during phototransduction (*13, 14*). Although immunogold electron microscopy has suggested peripherin-2 within incisures, the low labeling efficiency and sparse coverage inherent to this technique have precluded comprehensive protein mapping at the single-disc level (*15, 16*).

The connecting cilium bridges the inner and outer segments, functioning as both a modified transition zone and a selective protein trafficking conduit (*17–19*). This structure contains a 9+0 axoneme extending from the basal body, along with distal appendages (DAPs) and subdistal appendages (sDAPs), that anchor the cilium to cellular membranes (*2, 20–24*). Despite their structural importance, the nanoscale architecture of these components has been challenging to resolve in intact photoreceptors, owing to fundamental limitations of existing imaging approaches.

A fundamental incompatibility between resolution and molecular identification has constrained efforts to determine native disc molecular organization. To date, only electron microscopy has resolved individual photoreceptor discs, yielding virtually all current ultrastructural knowledge of disc architecture, yet it relies on non-specific structural contrast and cannot identify which proteins occupy specific regions (*2, 3*). Immunogold labeling attempts to bridge this gap, but suffers from low labeling efficiency (∼10–20% of available epitopes), steric hindrance from the large gold particles (10–30 nm), and sparse labeling that precludes comprehensive protein mapping at the single-disc level (*15, 16*).

In contrast, fluorescence microscopy achieves molecular specificity through immunolabeling, but cannot resolve structures separated by less than ∼200 nm—nearly sixfold larger than inter-disc spacing. Super-resolution techniques have improved this limit: stimulated emission depletion (STED) microscopy reaches 50–80 nm, and single-molecule localization microscopy (SMLM) attains 20–30 nm localization precision (*25, 26*). Yet even these advances remain insufficient to reliably resolve individual discs within the densely packed outer segment, where periodic structures repeat every 30–35 nm. This technical barrier leaves fundamental questions about native outer segment organization experimentally unresolved: how densely rhodopsin populates individual disc lamellae, whether peripherin-2 lines incisure membranes, and how the connecting cilium reorganizes structurally between inner and outer segments.

Expansion microscopy (ExM) addresses these limitations through physical specimen expansion within an isotropic hydrogel (*27–30*). In iterative ultrastructure expansion microscopy (iU-ExM), specimens undergo two sequential expansions for a total ∼20-fold magnification, achieving 10–20 nm effective resolution with antibody-based molecular identification (*31, 32*). Critically, the iterative strategy reduces effective antibody linkage error to 4–6 nm in original tissue dimensions, enabling accurate protein localization within structures separated by only 30–35 nm (see Materials and Methods) (*31, 33*).

Here, we employed iU-ExM to bridge the longstanding gap between electron microscopy and fluorescence-based imaging, enabling direct optical visualization of proteins within individual photoreceptor discs in intact tissue. Through 20-fold expansion, we visualized rhodopsin within individual discs, demonstrating that rhodopsin occupies approximately 92% of the disc lamellar domain in situ, and detected peripherin-2 within disc incisures. We further resolved connecting cilium architecture and established centriolar appendage dimensions as nanoscale reference values in photoreceptors. Application to the RCS rat model of retinitis pigmentosa reveals compartmentalized pathology: outer segment disc spacing increased 29% while inner segment centriolar architecture remained preserved, suggesting that early degeneration selectively disrupts membrane organization while sparing protein trafficking machinery (*34, 35*). These findings establish expansion microscopy as a powerful approach for interrogating photoreceptor organization in health and disease, with broad implications for understanding visual processing and the pathogenesis of inherited retinal degenerations.

## Results

### Protein Localization in Mammalian Photoreceptor Discs - Overall Organization of Expanded Retinal Tissue

To assess tissue preservation and structural integrity following expansion, we examined retinal sections at low magnification (*24, 31*). Retinal cell layers remained well-preserved throughout the expansion process. Combined rhodopsin and α/β-tubulin immunolabeling revealed complete photoreceptor architecture, with individual cilia visualized extending from the outer nuclear layer, through the inner segment, to the rod outer segment (Fig. 1A).

**Figure 1.**
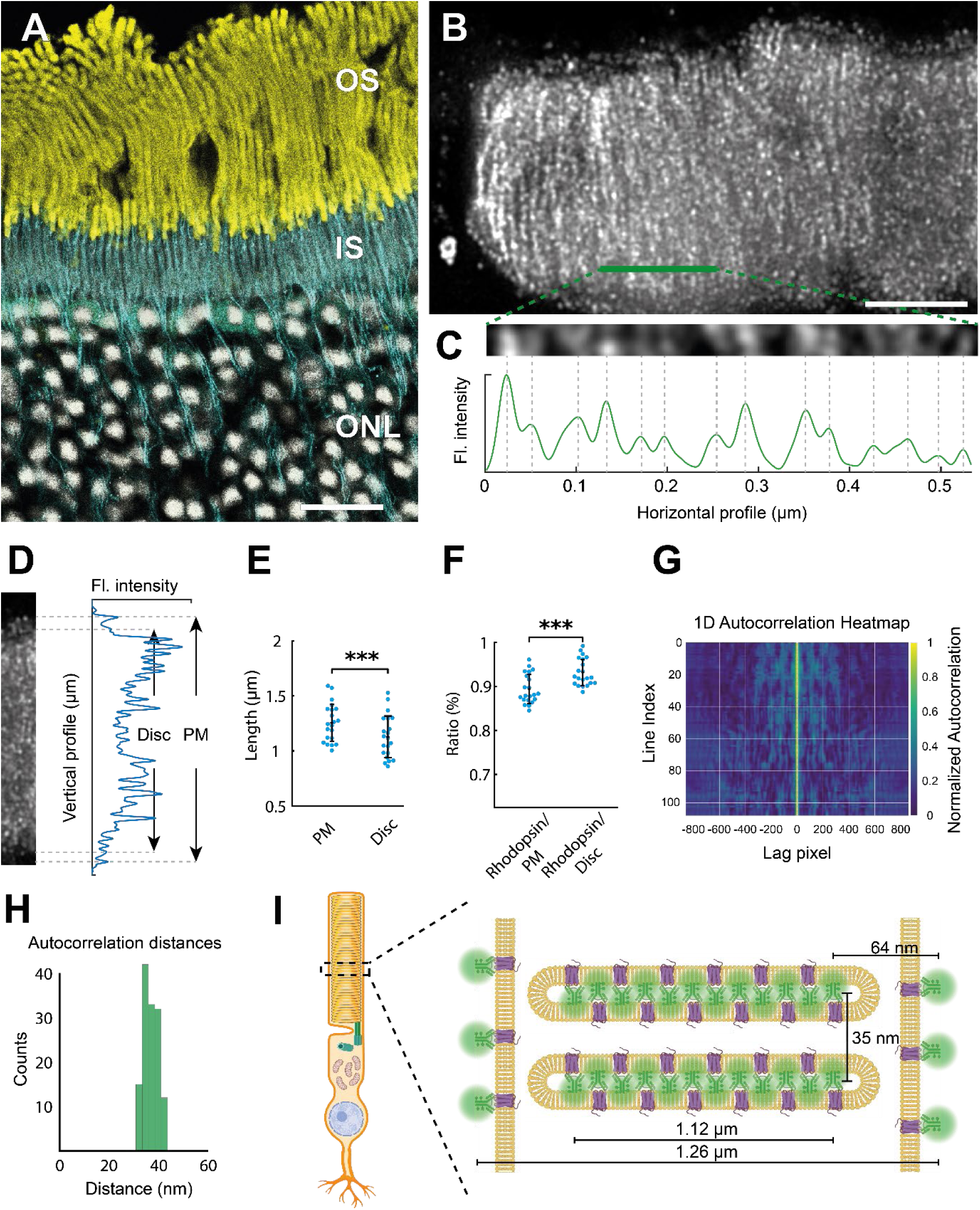
Rhodopsin visualization within individual photoreceptor discs reveals near-complete membrane occupancy. **A.** Low-magnification overview of 20-fold expanded rat retina showing preserved tissue architecture. Rhodopsin (yellow) identifies rod outer segments (OS), while α/β-tubulin immunolabeling (cyan) delineates cilia extending through the inner segments (IS). Nuclei (white) mark the outer nuclear layer (ONL). Scale bar, 20 µm. **B.** High-magnification view showing rhodopsin immunolabeling within a single photoreceptor outer segment reveals a highly ordered periodic pattern perpendicular to the longitudinal axis. Plasma membrane (PM) appears as a hollow cylinder surrounding the disc stack. Green outline indicates the region from which the intensity profile in C was derived. Scale bar, 500 nm. **C.** Horizontal intensity profile across multiple discs showing uniform periodicity. Individual peaks correspond to single discs. Direct peak-to-peak measurements yielded 35 ± 29 nm spacing (mean ± s.d.; CV = 83%); robust periodicity was therefore quantified by autocorrelation analysis (G–H). **D.** Vertical intensity profile along the radial axis showing rhodopsin-occupied disc diameter versus total outer segment diameter. Dashed lines indicate measurement boundaries. **E.** Quantification of radial extent of rhodopsin labeling (Disc) compared to total outer segment diameter (PM). Individual measurements shown as dots; center line, mean; error bars, s.d. PM: 1.26 ± 0.17 µm; Disc: 1.12 ± 0.19 µm; n = 42 profiles. P < 0.001, unpaired two-tailed t-test. **F.** Rhodopsin radial occupancy expressed as the ratio of rhodopsin-occupied diameter to plasma membrane diameter (Rhodopsin/PM) and to the disc lamellar domain (Rhodopsin/Disc). Individual measurements shown as dots; center line, mean; error bars, s.d. Rhodopsin/PM: 89 ± 3%; Rhodopsin/Disc: 92 ± 3%; n = 42 profiles. P < 0.001, unpaired two-tailed t-test. **G.** One-dimensional autocorrelation heatmap of rhodopsin signal across multiple line profiles (Line Index) confirms consistent disc periodicity. Central peak (lag = 0) and regular satellite peaks demonstrate uniform inter-disc spacing. Lag values (in pixels) were converted to physical distances after expansion factor correction for quantification in H. **H.** Histogram of autocorrelation-derived inter-disc distances showing a sharp peak at 35 ± 2.8 nm (mean ± s.d.), validating optical resolution of individual discs. N = 3417. **I.** Schematic representation of photoreceptor disc molecular organization based on measurements. Left: full rod photoreceptor with dashed box indicating the magnified region. Molecular components are color-coded: rhodopsin (purple) in disc lamellae; antibody point spread function (green); membrane (yellow). All measurements corrected for expansion factor.

Expansion factor was determined independently for each experiment by comparing photoreceptor nuclear cross-sectional area before and after expansion, yielding a mean linear expansion of 20.2 ± 2.65× (Fig. S1A; see Materials and Methods) (*31*).

Expansion accuracy was confirmed by two independent internal criteria: inter-disc periodicity matching established cryo-electron tomography values and preservation of ninefold radial symmetry in centriolar structures, both detailed in subsequent sections.

### Single-disc rhodopsin visualization reveals near-complete membrane occupancy

High-magnification rhodopsin immunolabeling revealed a highly ordered periodic pattern perpendicular to the outer segment axis, consistent with labeling of individual photoreceptor discs (Fig. 1B). Rhodopsin staining within discs was readily distinguished from plasma membrane signal, which exhibited a characteristic hollow cylindrical structure surrounding the disc stack.

To validate that this pattern represented individual discs, we first performed direct peak-to-peak measurements from intensity profiles, which yielded inter-disc spacing of 35 ± 29 nm (mean ± s.d.; CV = 83%; Fig. 1C, Fig. S1B). However, this approach suffered from high variability due to measurement noise inherent to individual profiles. To improve precision, we implemented one-dimensional autocorrelation analysis, which confirmed consistent disc periodicity of 35 ± 2.8 nm (CV = 8%; Fig. 1G–H). These values match established cryo-electron tomography measurements, validating optical resolution of individual discs and independently confirming expansion accuracy (*1, 3*). At 12 nm effective resolution, individual discs (35 nm spacing) are resolvable, but the two membranes forming each disc (separated by ∼4 nm intradiscal space) remain below resolution limits (Fig. S1C); each labeled band therefore represents a single disc (Fig. 1B).

Quantitative analysis along the disc radial axis revealed rhodopsin-occupied regions extending 1.12 ± 0.19 µm, compared to a total outer segment diameter of 1.26 ± 0.17 µm, corresponding to 89 ± 3% of outer segment cross-sectional diameter (Fig. 1D, E). To refine this estimate to the disc lamellar domain specifically, we measured the distance from outermost rhodopsin signal to plasma membrane (64 ± 15 nm) and subtracted plasma membrane-to-disc edge spacing (∼25 nm) and membrane thickness (∼7 nm) derived from electron microscopy literature (*2*). This yielded a rhodopsin-free zone of 32 ± 15 nm inward from the disc rim. Accounting for this peripheral exclusion zone, rhodopsin occupies approximately 92 ± 3% of the disc lamellar domain (Fig. 1F, I; Fig. S1D). This estimate incorporates antibody-defined radial boundaries, internally validated isotropic expansion, and literature-based rim geometry; coverage may be modestly influenced by probe size and labeling geometry.

This dense in situ packing substantially exceeds the approximately 50% membrane area fraction reported in atomic force microscopy studies of isolated disc membranes, while aligning with biochemical estimates that rhodopsin constitutes ∼90% of outer segment membrane protein mass (*6, 36*). The methodological basis for this divergence is examined in the Discussion.

### Peripherin-2 localizes within disc incisures

Peripherin-2 immunolabeling revealed the characteristic hollow cylindrical structure of photoreceptor outer segments (Fig. 2A) (*9–11*). Orthogonal views resolved individual disc morphology and, notably, peripherin-2 signal extending inward from the disc rim in a pattern consistent with incisure localization, extending previous genetic and immunogold evidence to direct optical protein visualization (Fig. 2B) (*37*).

**Figure 2.**
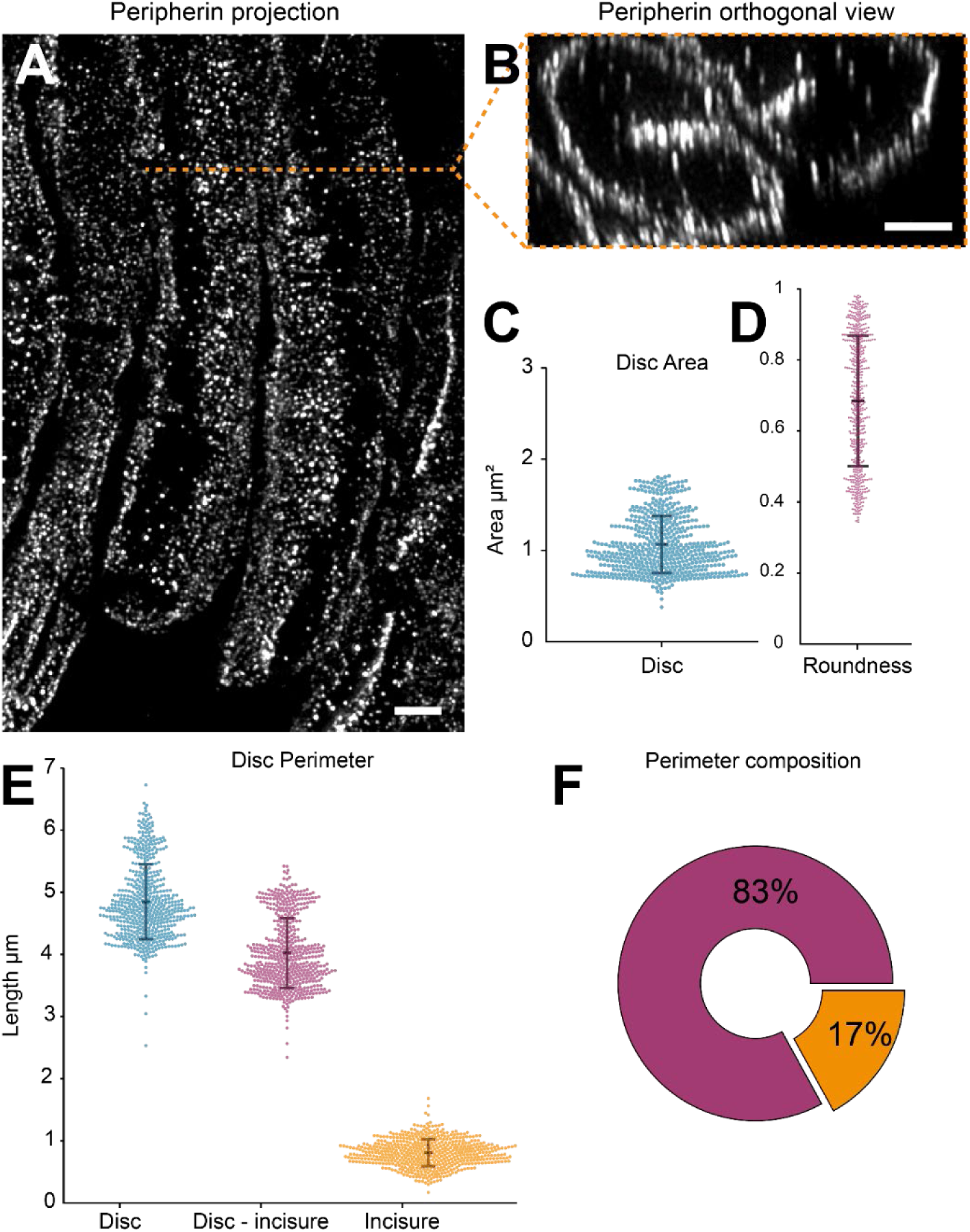
Peripherin-2 localizes within disc incisures. **A.** Maximum intensity projection of peripherin-2 immunolabeling showing the characteristic hollow cylindrical structure of multiple photoreceptor outer segments. Individual rod outer segments display continuous peripherin-2 signal along their length, delineating disc rim boundaries. Scale bar, 0.5 µm. **B.** Orthogonal view of peripherin-2-labeled discs reveals detailed disc morphology. Projection showing stacked disc rims, demonstrating disc roundness with peripherin-2 signal extending previous genetic and ultrastructural evidence to direct optical protein visualization within these structures. Orange dashed line indicates the plane of orthogonal reconstruction. Scale bar, 0.5 µm. **C.** Quantification of disc area measured from peripherin-2-labeled en face views. Individual discs shown as dots; center line, mean; error bars, s.d. n = 547. **D.** Disc roundness index distribution. Values approaching 1.0 indicate perfect roundness. Individual discs shown as dots; center line, mean; error bars, s.d. n = 547. **E.** Disc perimeter measurements showing total perimeter (Disc) (blue), perimeter excluding incisures (Disc − incisure) (magenta), and incisure membrane perimeter (Incisure) (orange). Individual discs shown as dots; center line, mean; error bars, s.d. n = 547. **F.** Donut chart showing mean proportional contributions of incisures (17%) (orange) and external rim (83%) (purple) to total disc perimeter. n = 547. All measurements corrected for expansion factor.

The ability to resolve peripherin-2 at the single-disc level enabled quantitative characterization of disc geometry and incisure organization, establishing baseline values for subsequent disease model comparison. Overall, discs exhibited near-circular profiles (roundness index 0.69 ± 0.18) with a mean area of 1.12 ± 0.33 µm² and total perimeter of 4.86 ± 0.60 µm (Fig. 2C–E), maintaining uniform geometry along the photoreceptor length.

Within this perimeter, incisures contributed 0.82 ± 0.11 µm (17 ± 0.4% of total perimeter; Fig. 2E–F), representing the membrane path length traced along both arms of each fold, as the lipid bilayer lines both sides of the cleft. The corresponding incisure depth, the linear distance from disc rim to the deepest point, measured 0.41 ± 0.055 µm. The remaining external rim perimeter measured 4.03 ± 0.56 µm. This incisure-to-perimeter ratio remained consistent across all measured discs, indicating that the proportion of rim allocated to incisures is tightly regulated during disc morphogenesis.

### Connecting cilium undergoes defined morphological transformation at the outer segment boundary

The connecting cilium constitutes a critical structural element supporting outer segment elongation and maintenance of photoreceptor architecture (*17, 18*). To characterize the structural transition between inner and outer segments, we performed three-dimensional reconstruction of individual microtubule filaments, with particular focus on morphological changes at the connecting cilium (Fig. 3A). These measurements establish baseline ciliary architecture for subsequent comparison with the RCS disease model.

**Figure 3.**
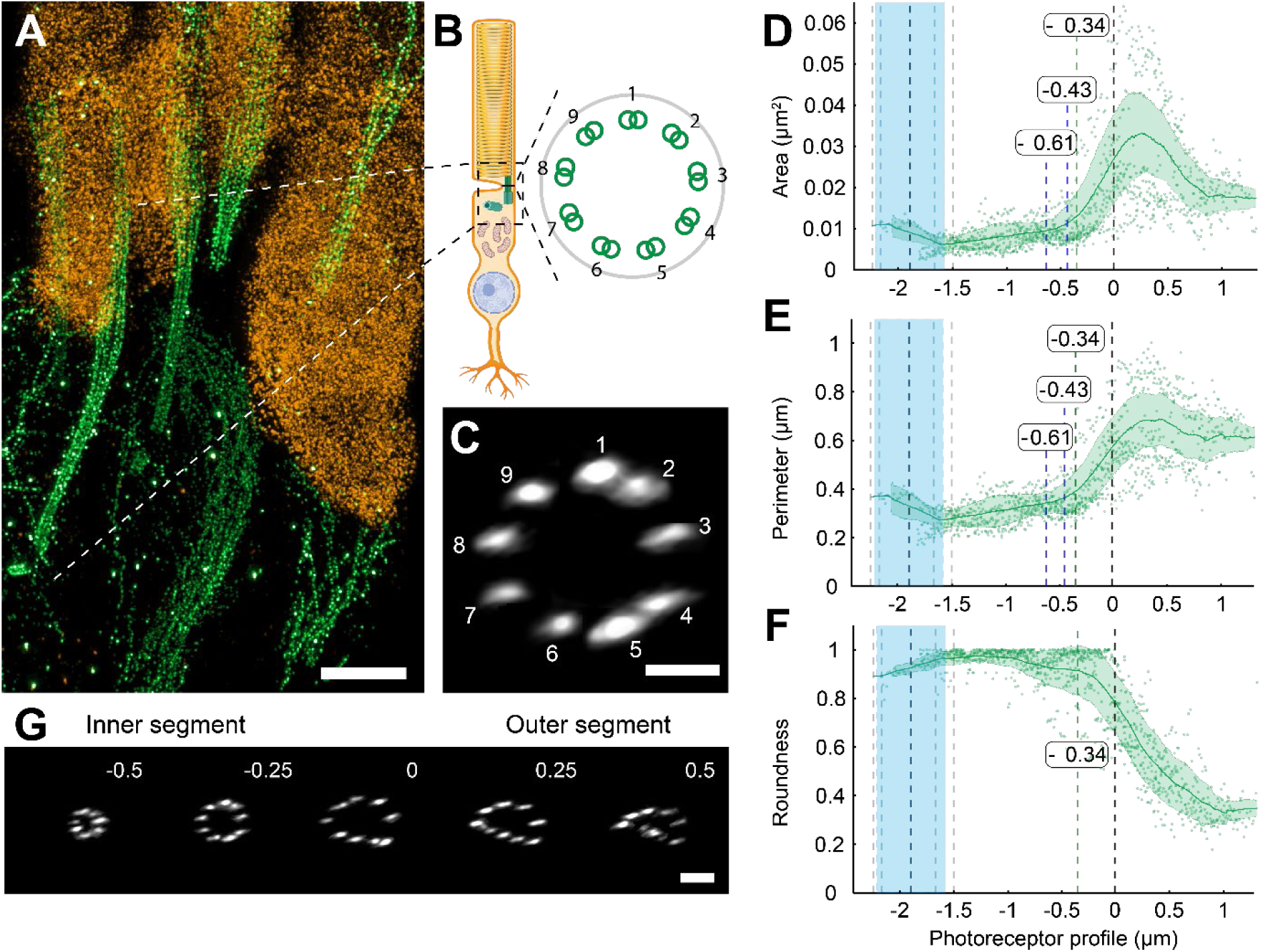
Three-dimensional reconstruction reveals connecting cilium morphological transformation during inner-to-outer segment transition. **A.** Composite image showing photoreceptor architecture with tubulin labeling (green) delineating the connecting cilium from the basal body through the inner segment (IS) to the outer segment (OS). Rhodopsin (orange) marks the disc-containing outer segment. White dashed lines indicate the position of cross-sectional analysis. Scale bar, 1 µm. **B.** Schematic of photoreceptor architecture illustrating the position of the connecting cilium and the 9+0 axonemal arrangement. Inset: nine microtubule doublets arranged in characteristic radial symmetry. **C.** Cross-sectional view of the connecting cilium showing nine individually resolved microtubule doublets (numbered 1–9) arranged in the 9+0 configuration, confirming preservation of ninefold radial symmetry after expansion. Scale bar, 100 nm. **D.** Ciliary cross-sectional area as a function of axial position relative to the inner-outer segment junction (0 µm). Blue shaded region indicates basal body position. The cilium maintains compact geometry until morphological transformation initiates proximal to the outer segment. n = 8 cilia. **E.** Ciliary perimeter as a function of axial position, following the same spatial progression as area expansion. n = 8 cilia. **F.** Roundness as a function of axial position. High roundness is maintained throughout the inner segment, with progressive decline through the transition zone, quantifying the transition from circular to asymmetric expanded configuration. n = 8 cilia. **G.** Serial cross-sectional images of the connecting cilium at defined axial positions (−0.5, −0.25, 0, +0.25, +0.5 µm relative to the IS/OS junction) illustrating the progressive transition from the compact nine-doublet axonemal arrangement in the inner segment to the expanded, asymmetric microtubule configuration in the outer segment. Scale bar, 200 nm. All measurements corrected for expansion factor.

Within the inner segment, we resolved the characteristic nine microtubule triplets of the basal body and nine doublets comprising the 9+0 axoneme (Fig. 3B, C) (*2, 17*). Alignment of ciliary traces at the outer segment entry point positioned the basal body at 1.87 ± 0.36 µm proximal to the outer segment boundary.

Throughout the inner segment, the cilium maintained compact, circular geometry (roundness >0.9) with cross-sectional area of 6,190 ± 1,360 nm² and perimeter of 276 ± 31 nm (Fig. 3D–E). This configuration persisted until 0.61 µm proximal to the outer segment, where the area curve first departed from baseline. Derivative analysis (see Materials and Methods) identified two landmarks in this transformation: an onset at 0.43 µm, where the rate of area increase reached its maximum (area: 11,000 ± 4,000 nm²; perimeter: 363 ± 65 nm), and a peak at 0.34 µm, where morphological change was greatest (area: 33,000 ± 9,000 nm²; perimeter: 693 ± 109 nm; Fig. 3D–E, Fig. S2).

This transformation was accompanied by progressive loss of circular symmetry. Roundness declined from 0.9–1.0 within the inner segment to 0.32 in the outer segment (measured 1 µm distal to the junction; Fig. 3F). Serial cross-sections at defined axial positions (−0.5, −0.25, 0, +0.25, +0.5 µm relative to the junction) illustrate this progression from compact nine-doublet arrangement to expanded, asymmetric microtubule configuration (Fig. 3G). Within the outer segment proper, 9+0 axonemal symmetry was disrupted, yielding irregular microtubule organization.

### NHS-ester labeling resolves centriolar appendage architecture

Centriolar appendages mediate ciliary vesicle docking (DAPs) and cytoplasmic microtubule anchoring (sDAPs), yet their nanometer-scale dimensions and close spatial proximity (∼100 nm separation) have restricted their characterization in photoreceptors to electron microscopy (*20, 21, 23*). To resolve these structures optically, we employed NHS-ester protein density mapping, a chemical labeling approach that targets primary amine residues and generates comprehensive protein density maps analogous to electron microscopy contrast while retaining fluorescence-based detection (*30, 31*) (Fig. 4A).

**Figure 4.**
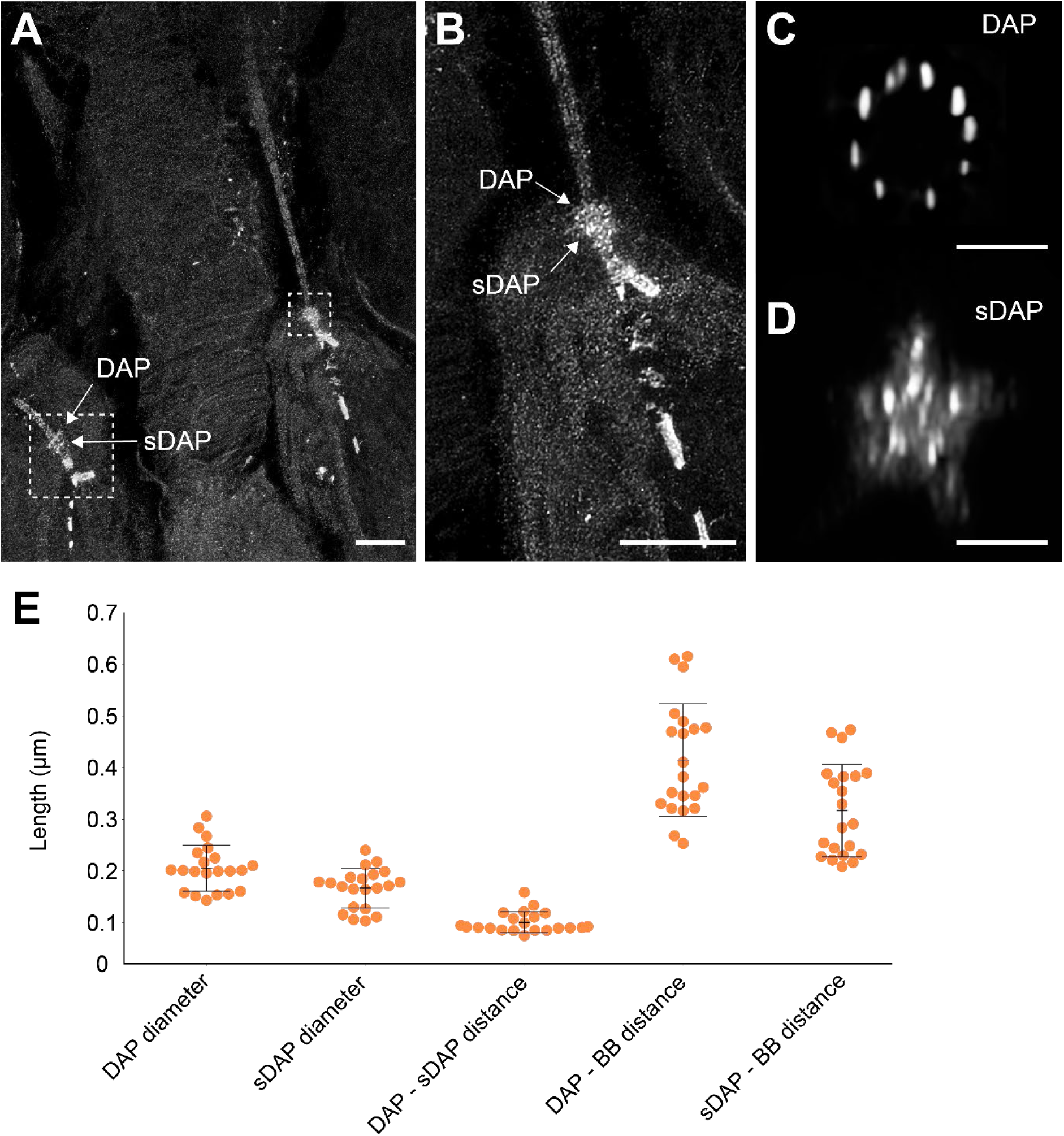
NHS-ester protein density mapping resolves centriolar appendage architecture in photoreceptors. **A.** Maximum intensity projection of NHS-ester labeling providing comprehensive ultrastructural visualization of photoreceptor architecture. This chemical labeling approach targets primary amine residues, generating protein density maps. Disc membranes, connecting cilium, and centriolar appendages are simultaneously resolved. Dashed boxes indicate centriolar appendages: distal appendages (DAP) and subdistal appendages (sDAP) are indicated. Scale bar, 0.5 µm. **B.** Magnified view of the basal body region showing the spatial relationship between DAP and sDAP structures. The ninefold radial symmetry of both appendage types is visible, oriented perpendicular to the ciliary axis at the inner segment periphery. Scale bar, 0.5 µm. **C.** En face view of distal appendages revealing characteristic ninefold radial symmetry. Nine discrete protein density foci correspond to DAP blades mediating ciliary vesicle docking. Scale bar, 100 nm. **D.** En face view of subdistal appendages displaying radial symmetry. These structures function in cytoplasmic microtubule anchoring. Scale bar, 100 nm. **E.** Quantification of centriolar appendage dimensions: DAP diameter, sDAP diameter, DAP–sDAP axial spacing, and DAP and sDAP distances from the basal body (BB). Individual measurements shown as dots; center line, mean; error bars, s.d. DAP: 205 ± 44 nm; sDAP: 167 ± 38 nm; spacing: 101 ± 20 nm; DAP-BB: 415 ± 109 nm; sDAP-BB: 317 ± 89 nm; all n = 21. All measurements corrected for expansion factor.

This approach simultaneously resolved multiple photoreceptor compartments, including the connecting cilium with nine axonemal subunits, disc membranes, and mitochondrial networks (Fig. 4A, B). Critically, both DAPs and sDAPs were visualized, each displaying characteristic ninefold radial symmetry oriented perpendicular to the ciliary axis at the inner segment periphery (Fig. 4C, D) (*20, 21, 23*).

To establish dimensional reference values for the inner-outer segment transition zone, we performed morphometric quantification. DAP diameter measured 205 ± 44 nm and sDAP diameter 167 ± 38 nm. The two structures were separated by 101 ± 20 nm along the ciliary axis, with the DAP ring positioned at 415 ± 109 nm and the sDAP ring at 317 ± 89 nm from the basal body (Fig. 4E). This spatial arrangement places the DAP ring at the distal boundary of the basal body, marking the origin of the connecting cilium toward the outer segment. These baseline values, together with the disc and ciliary morphometry described above, provide the wild-type reference framework for detecting structural alterations in the RCS disease model.

### Compartmentalized pathology in the RCS rat model of retinitis pigmentosa

To investigate whether iU-ExM could detect structural alterations in retinal disease, we examined photoreceptors from Royal College of Surgeons (RCS) rats at postnatal day 30 (P30). The RCS rat harbors a mutation in the Mertk gene that renders retinal pigment epithelium cells unable to phagocytose shed outer segment material, leading to debris accumulation and progressive photoreceptor degeneration (*34, 35*). At P30, outer segment disruption and subretinal debris accumulation are evident, though substantial photoreceptor loss has not yet occurred, making this time point well suited for interrogating early-to-moderate pathology (*38*).

### Increased inter-disc spacing despite preserved disc dimensions in RCS photoreceptors

At P30, the majority of RCS outer segments exhibited advanced disruption incompatible with disc-resolved analysis. We identified and analyzed the subset of outer segments retaining sufficient structural integrity for autocorrelation-based periodicity measurement (Fig. 5A; see Materials and Methods). These findings therefore characterize disc organization in the small fraction of photoreceptors that have not yet undergone severe structural collapse, representing the earliest detectable stage of pathology rather than the predominant disease phenotype at this time point. Despite visible structural damage including membrane discontinuities, protein distribution patterns in analyzable segments resembled those of wild-type photoreceptors (expansion factor determination followed the same nuclear area-based approach as wild-type; Fig. S3A).

**Figure 5.**
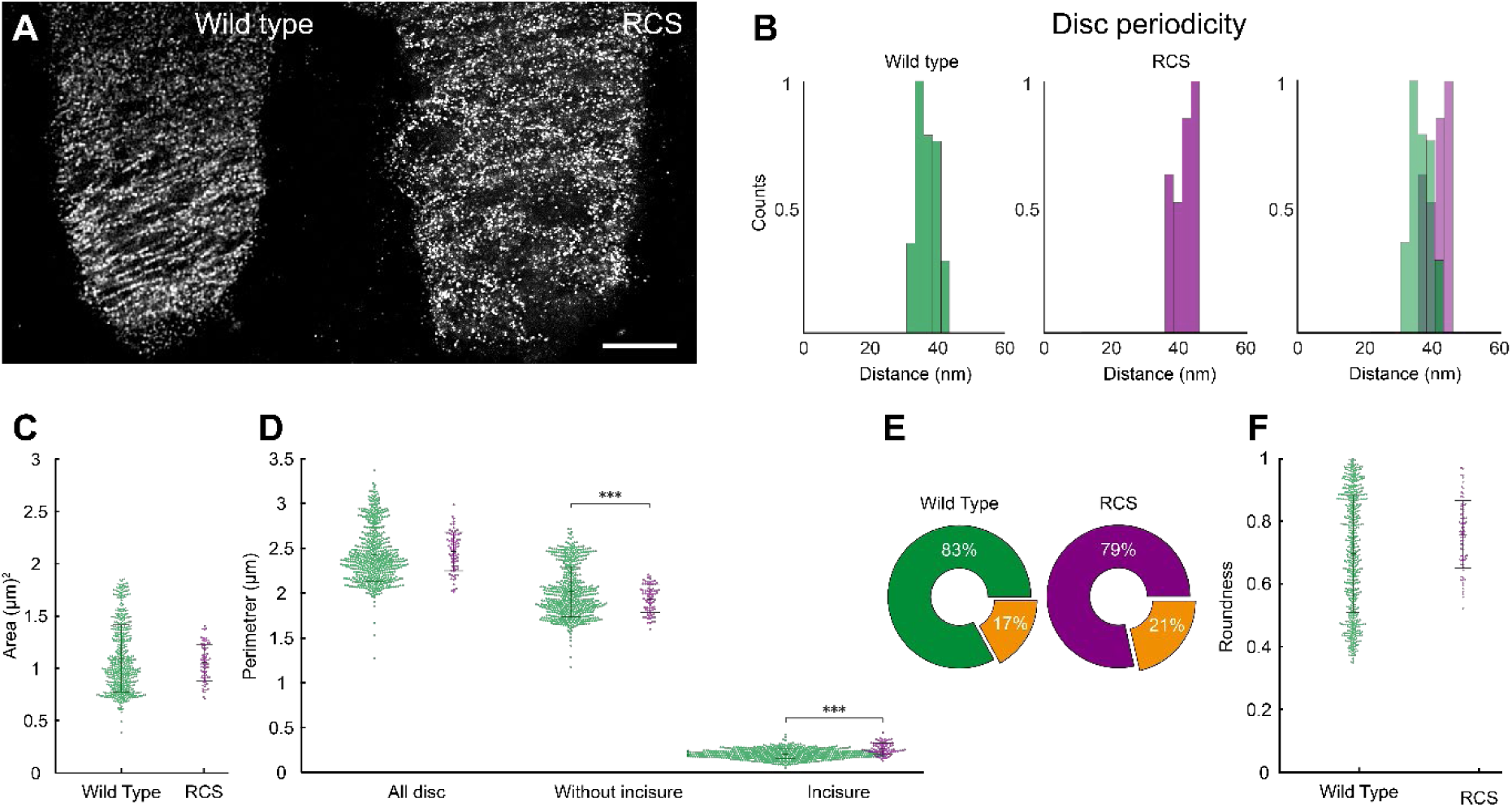
Nanoscale disc pathology in the RCS rat model of retinitis pigmentosa. **A.** Rhodopsin immunolabeling of photoreceptor outer segments from wild-type (left) and RCS (right) rats. Wild-type photoreceptors display organized disc architecture with uniform spacing, while RCS photoreceptors exhibit disrupted disc organization with visible structural irregularities despite preserved overall protein distribution patterns. Scale bar, 500 nm. **B.** Histograms of autocorrelation-derived inter-disc spacing. Wild-type (left, green) shows a sharp peak at 35 nm; RCS (centre, purple) displays broader peaks shifted toward longer periodicities (∼45 nm); overlaid comparison (right) demonstrates a 29% increase in inter-disc spacing. **C.** Disc area comparison between genotypes. Individual discs shown as dots; center line, mean; error bars, s.d. WT (green), RCS (purple). n = 547 (WT), n = 77 (RCS) discs. P = 0.1, unpaired two-tailed t-test; not significant. **D.** Disc perimeter analysis showing total perimeter, perimeter excluding incisures, and incisure membrane perimeter for WT (green) and RCS (purple). Individual discs shown as dots; center line, mean; error bars, s.d. n = 547 (WT), n = 77 (RCS) discs. Total perimeter, P = 0.2; perimeter excluding incisures, P < 0.001; incisure perimeter, P < 0.001; unpaired two-tailed t-tests. **E.** Donut charts showing mean proportional contributions of external rim (green and purple) and incisures (orange) to total disc perimeter in wild-type (left) and RCS (right). n = 547 (WT), n = 77 (RCS) discs. **F.** Disc roundness comparison. Individual discs shown as dots; center line, mean; error bars, s.d. WT (green), RCS (purple). n = 547 (WT), n = 77 (RCS) discs. P = 0.73, unpaired two-tailed t-test. All measurements corrected for expansion factor.

Autocorrelation analysis of inter-disc spacing revealed sharp peaks at 35 nm in wild-type photoreceptors, whereas RCS samples exhibited broader peaks shifted toward longer periodicities (Fig. 5B). Direct measurement confirmed increased inter-disc spacing in RCS segments: 43.8 ± 4.9 nm compared to 35 ± 2.8 nm in wild-type (Fig. 5B, Fig. S3B), representing a ∼29% increase. Importantly, disc area and perimeter showed no significant differences between genotypes (see below; Fig. 5C), arguing against differential expansion of outer segment compartments as an explanation for the increased spacing.

Despite this altered spacing, peripherin-2 immunolabeling of en face disc profiles revealed no significant differences between genotypes in total disc area (WT: 1.12 ± 0.33 µm²; RCS: 1.07 ± 0.1 µm²; P = 0.1) or total disc perimeter (WT: 4.86 ± 0.60 µm; RCS: 4.93 ± 0.43 µm; P = 0.2) (Fig. 5C-D). Disc roundness was similarly maintained (WT: 0.695 ± 0.186; RCS: 0.757 ± 0.108; P = 0.73), confirming that gross disc geometry remains intact in the disease model (Fig. 5F).

Within this preserved geometry, incisure organization was also retained. Incisure membrane perimeter was modestly increased in RCS discs (1.06 ± 0.108 µm vs. 0.82 ± 0.11 µm in wild-type), corresponding to incisure depths of 0.530 ± 0.108 µm and 0.41 ± 0.055 µm, respectively (Fig. 5D). This corresponded to incisure contributions of 21 ± 0.9% versus 17 ± 0.4% of total disc perimeter (Fig. 5E). The peripherin-2-excluded disc perimeter, i.e. the disc rim without incisures, showed comparable values between groups (Fig. 5D). Overall, these measurements indicate that disc morphogenesis remains fundamentally intact in RCS photoreceptors, with alterations confined to inter-disc spacing rather than disc structure itself.

### Connecting cilium transformation zone shifts proximally in RCS photoreceptors

Having established that disc spacing is altered while disc dimensions are preserved, we next asked whether pathology extends to the connecting cilium, which bridges the affected outer segment and the inner segment. Morphometric analysis of ciliary cross-sections at 50 nm axial intervals revealed that the transition zone, where the compact axoneme begins expanding into the outer segment configuration, occurred closer to the outer segment in RCS photoreceptors (0.36 µm) compared to wild-type (0.43 µm; P < 0.03; Fig. 6B-C, Fig. S4). This ∼70 nm proximal shift was accompanied by a reduced basal body-to-outer segment distance (RCS: 1.71 ± 0.33 µm; WT: 1.87 ± 0.36 µm; Fig. 6A). Once initiated, the morphological transformation followed a similar trajectory to wild-type, with coordinated increases in area and perimeter and progressive loss of roundness (Fig. 6B-C). However, RCS photoreceptors reached slightly smaller absolute dimensions throughout the transition zone, suggesting the same reorganization program operating at reduced magnitude (Fig. 6C).

**Figure 6.**
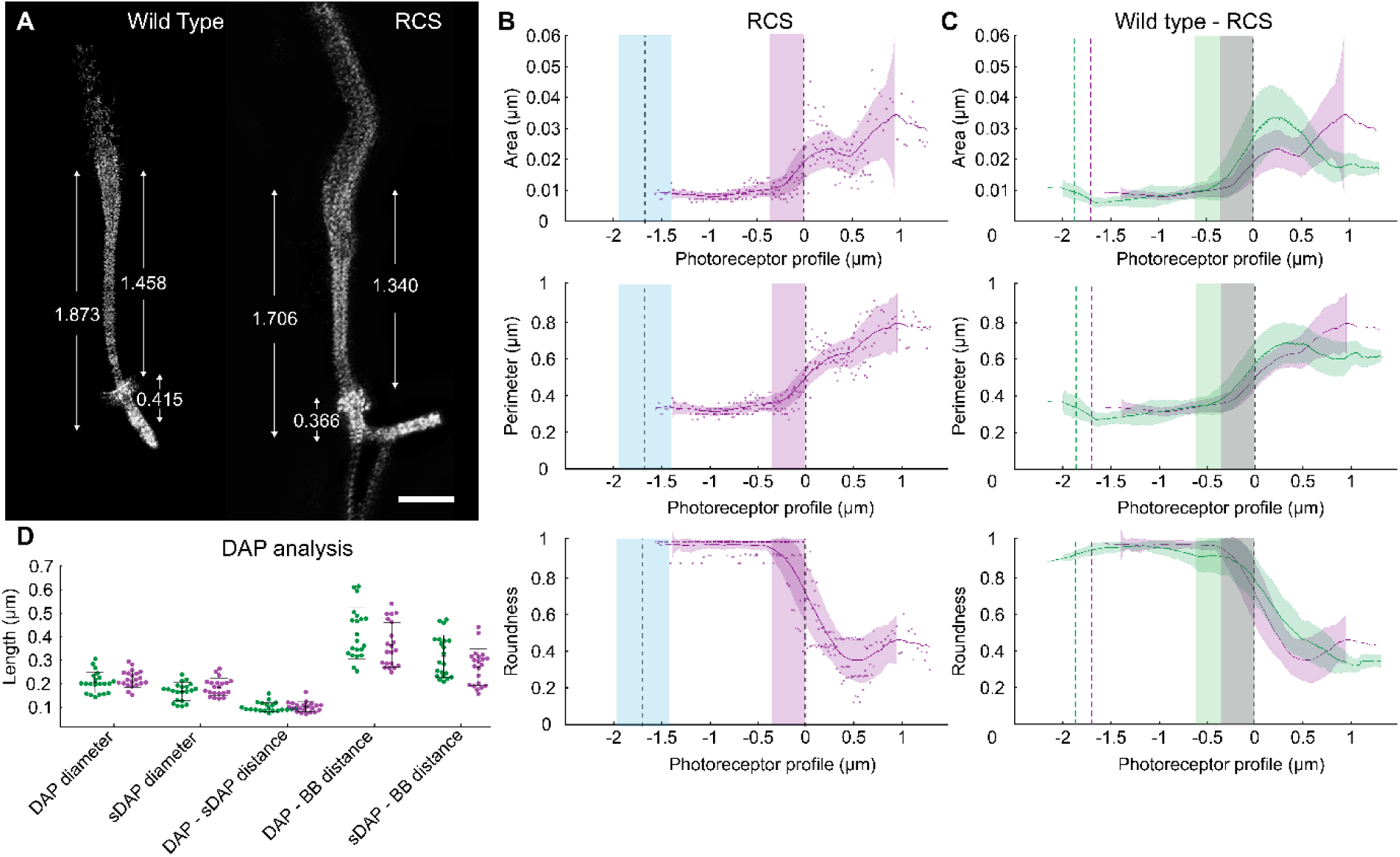
Connecting cilium morphology and centriolar appendage preservation in the RCS disease model. **A.** Representative NHS-ester-labeled connecting cilia from wild-type (left) and RCS (right) photoreceptors with dimensional annotations in (nm). Overall ciliary architecture is preserved in RCS despite altered positioning relative to outer segment. Scale bars, 500 nm. **B.** Morphometric analysis of RCS connecting cilium transformation. Area (top), perimeter (middle), and roundness (bottom) plotted as a function of axial position relative to the inner-outer segment junction (0 µm). Individual measurements shown as dots; solid line, smoothed mean; shaded band, s.d. Blue shaded region indicates basal body position; purple shaded region marks the morphological transformation zone. n = 6 cilia. **C.** Direct comparison of wild-type (green) and RCS (purple) morphometric profiles. Area (top), perimeter (middle), and roundness (bottom). While absolute values differ, the fundamental pattern of ciliary transformation is preserved between genotypes. The transition zone initiates at 0.36 µm in RCS (compared to 0.43 µm in WT), representing a ∼70 nm proximal shift. n = 8 (WT), n = 6 (RCS) connecting cilia. **D.** Centriolar appendage architecture comparison between wild-type (green) and RCS (purple). Measurements include DAP diameter, sDAP diameter, inter-appendage spacing, and distances from basal body. Individual measurements shown as dots; center line, mean; error bars, s.d. n = 21 (WT), n = 21 (RCS) appendage sets. DAP diameter, P = 0.22; sDAP diameter, P = 0.10; DAP-sDAP spacing, P = 0.76; DAP-BB distance, P = 0.16; sDAP-BB distance, P = 0.11; unpaired two-tailed t-tests. All measurements corrected for expansion factor.

### Centriolar appendage architecture is preserved despite outer segment pathology

To determine whether structural alterations propagate further into the inner segment, we examined DAP and sDAP architecture. Morphometric analysis revealed no significant differences between genotypes in DAP diameter (WT: 205 ± 44 nm; RCS: 221 ± 35 nm), sDAP diameter (WT: 167 ± 38 nm; RCS: 186 ± 37 nm), inter-appendage spacing (WT: 101 ± 20 nm; RCS: 102 ± 21 nm), or positions relative to the basal body of DAPs (WT: 415 ± 109 nm; RCS: 366 ± 94 nm) and sDAPs (WT: 317 ± 89 nm; RCS: 271 ± 77 nm) (Fig. 6D). This preservation indicates that centriolar appendages, and the protein trafficking infrastructure they support, remain resilient to the early degenerative changes affecting the outer segment.

## Discussion

Despite six decades of investigation, the molecular organization of photoreceptor discs has eluded direct optical imaging at the single-disc level. This study demonstrates that iterative ultrastructure expansion microscopy enables direct optical visualization of proteins within individual photoreceptor discs, integrating molecular identification with ultrastructural context at 12 nm effective resolution. The highly ordered, periodic rhodopsin labeling pattern with 35 ± 2.8 nm inter-disc spacing, precisely matching cryo-electron tomography measurements (*1, 3*), confirms that individual discs are resolved and that the proteins they contain are directly visualized. Rather than merely improving resolution of previously observable structures, this approach integrates molecular identification with ultrastructural context. It enables quantification of protein distributions within individual membrane compartments, correlation of molecular organization with disc position along the outer segment, and detection of early organizational changes in disease models before gross structural degeneration becomes evident.

Our quantitative analysis revealed that rhodopsin occupies 92 ± 3% of the disc lamellar domain in situ, substantially exceeding the approximately 50% area fraction reported in atomic force microscopy studies of isolated disc membranes (*8, 36, 39*). However, these metrics are not directly comparable: our radial occupancy measurement quantifies the lateral extent of antibody-accessible rhodopsin within intact tissue, whereas AFM area fractions derive from topographic imaging of isolated, substrate-adsorbed membranes. Assuming circular disc geometry, the corresponding area fractions would be ∼79% (PM boundary) and ∼85% (lamellar domain), still substantially exceeding AFM estimates.

Furthermore, our coverage calculation assumes isotropic radial expansion and employs literature-derived values for plasma membrane-to-disc edge spacing. Nonetheless, the magnitude of the divergence suggests that AFM preparations underestimate native rhodopsin density. Isolation of disc membranes requires osmotic lysis, detergent treatment, and substrate adsorption, each step potentially disrupting native protein organization (*8*); loosely associated rhodopsin molecules may dissociate during extraction, and flattening curved membranes onto mica substrates may alter packing geometry. The concordance of our in situ estimate with biochemical data indicating that rhodopsin constitutes approximately 90% of outer segment membrane protein mass supports this interpretation (*6*). Moreover, the SDS denaturation inherent to iU-ExM processing unfolds proteins and maximizes epitope accessibility throughout the lamellar domain, reducing the likelihood that our estimate is limited by systematic epitope masking near specialized membrane regions.

This near-complete occupancy has functional implications for phototransduction: dense rhodopsin packing maximizes photon capture probability, critical for the single-photon sensitivity that defines rod function, while potentially constraining lateral diffusion of activated rhodopsin and downstream signaling molecules. The 32 nm rhodopsin-free peripheral zone corresponds to the peripherin-2/ROM1-enriched rim domain, indicating functional compartmentalization between the phototransduction-active disc lamella and the structurally specialized rim. This revised estimate also has practical implications: therapeutic strategies aimed at modulating rhodopsin expression, including gene augmentation and antisense oligonucleotide approaches, may require recalibration of target protein densities to match native disc organization.

Direct visualization of peripherin-2 within disc incisures extends previous observations to comprehensive molecular mapping at the single-disc level. Immunogold electron microscopy has localized peripherin-2 to disc rims, including regions corresponding to incisures, but the inherent limitations of this approach, sparse labeling efficiency (10–20% of epitopes), large probe size, and inability to resolve protein distribution within individual structural features, precluded comprehensive characterization of peripherin-2 organization within incisures (*10, 11*). Genetic studies provided complementary evidence: peripherin-2 deficiency suppresses incisure formation, while overexpression produces elongated incisures, implying a structural role without directly demonstrating molecular presence (*9, 12, 37*). Our data show that peripherin-2 lines incisure membranes continuously along both arms of each fold, confirming that incisures are stabilized by the same tetraspanin complexes that fortify external disc rims. The uniform incisure-to-perimeter ratio (17 ± 0.4%) is consistent with tight stoichiometric control during disc morphogenesis, reflecting the molar relationship between rhodopsin and peripherin-2/ROM1 expression (*37*). These morphometric parameters provide quantitative constraints for computational models of phototransduction cascade kinetics, where incisure depth and distribution influence diffusional coupling across the disc surface.

Three-dimensional reconstruction of tubulin-labeled connecting cilia revealed a defined morphological transformation between inner and outer segments. Within the inner segment, the 9+0 axoneme retains compact, circular geometry (roundness >0.9) until ∼430 nm proximal to the outer segment boundary. Then, cross-sectional area expands 5.3-fold with progressive loss of circular symmetry, accommodating the lateral membrane outgrowth required for nascent disc formation (Fig. 3D-E). Within the outer segment, the axonemal symmetry becomes disrupted, with microtubule doublets splaying into an irregular arrangement alongside the maturing disc stack. This transformation correlates spatially with the disc evagination zone, suggesting functional coupling between axonemal reorganization and membrane morphogenesis. The tight circular configuration maintained in the inner segment reflects the structural constraints imposed by the transition zone, where Y-links and the ciliary necklace establish the diffusion barrier maintaining outer segment protein compartmentalization (*2, 24*).

Complementing antibody-based tubulin imaging, NHS-ester protein density mapping resolved centriolar appendage architecture at the inner-outer segment junction (Fig. 4). DAPs mediate ciliary vesicle docking essential for ciliogenesis, while sDAPs anchor cytoplasmic microtubules; despite their critical roles, these structures have been characterized in photoreceptors exclusively through electron microscopy due to their nanometer-scale dimensions and close spatial proximity. Our measurements established DAP diameter of 205 ± 44 nm and sDAP diameter of 167 ± 38 nm, with ninefold radial symmetry clearly resolved, and inter-appendage spacing of 101 ± 20 nm positioning these structures at 415 ± 109 nm (DAP) and 317 ± 89 nm (sDAP) from the basal body (Fig. 4E). These values are consistent with super-resolution measurements in cultured cells, establishing optical reference values for centriolar appendages in photoreceptors that can now be interrogated for changes in disease models or developmental contexts (*2, 19, 20, 23, 32*).

To test the capacity of iU-ExM to detect pathological changes, we examined RCS photoreceptors at P30, a time point representing early-to-moderate pathology when outer segment disruption is evident but substantial photoreceptor loss has not yet occurred (*38*). Previous longitudinal studies established that early pathology in this model is compartmentalized, with outer segment degeneration progressing while inner retinal architecture remains stable (*38*). Our nanoscale analysis provides molecular-level support for this model and extends it to the subcellular architecture of individual photoreceptors.

Within the outer segment, inter-disc spacing increased ∼29% (Fig. 5B), likely reflecting altered cytoplasmic composition or osmotic imbalance secondary to debris accumulation. Autocorrelation analysis revealed broader peaks in RCS compared to the sharp periodicity in wild-type, indicating increased variability in disc organization. Preserved disc dimensions and centriolar architecture between genotypes argue against differential expansion artifacts, though we cannot exclude that local changes in crosslinking density within degenerated outer segments modestly influence spacing estimates. Despite this spacing alteration, disc dimensions, roundness, and incisure organization were all preserved (Fig. 5C-F), with a modest increase in incisure contribution to total perimeter (21% vs. 17%; Fig. 5E), indicating that the fundamental disc formation machinery continues to function normally even as the inter-disc environment deteriorates. These findings suggest that disc morphogenesis is not impaired in early RCS pathology; rather, the alterations reflect secondary consequences of impaired debris clearance on the environment surrounding otherwise normally formed discs.

Extending the spatial analysis inward, the connecting cilium exhibited a ∼70 nm proximal shift of the transformation zone (Fig. 6B-C). This alteration may reflect mechanical compression from accumulated debris at the outer segment apex or an adaptive response to altered outer segment length. Once initiated, however, the morphological transformation followed a trajectory similar to wild-type, though at reduced magnitude, indicating that the fundamental process of cilium-to-outer segment reorganization remains intact despite altered spatial positioning. Further into the inner segment, DAP and sDAP architecture showed no significant differences between genotypes (Fig. 6D). The preservation of these structures, which mediate ciliary vesicle docking and cytoplasmic microtubule anchoring, indicates that the protein trafficking infrastructure at the basal body is resilient to the early degenerative changes affecting the outer segment.

This selective vulnerability pattern, in which outer segment structures are disrupted while inner segment centriolar architecture remains stable, has mechanistic and therapeutic implications. The Mertk mutation generates pathology at the outer segment-RPE interface through debris accumulation, and our data suggest that this insult propagates inward, altering disc spacing and transition zone positioning, but does not compromise the inner segment structures that constitute the protein delivery gateway. The resilience of centriolar appendages and basal body architecture indicates that photoreceptors retain the capacity for protein trafficking even as their destination compartment degenerates, consistent with the stable inner retinal architecture observed during early RCS degeneration (*38*).

This compartmentalization defines a potential therapeutic window. Interventions targeting debris clearance or outer segment stabilization could prove effective precisely because the inner segment trafficking machinery remains functional. Gene therapy approaches restoring Mertk function, RPE transplantation strategies, or pharmacological interventions promoting alternative phagocytic pathways could potentially halt or reverse outer segment degeneration while the protein delivery infrastructure remains intact. By resolving these parameters at the single-disc level, our approach provides nanoscale biomarkers, including inter-disc spacing, transition zone position, and centriolar appendage dimensions, for assessing treatment efficacy before gross structural degeneration becomes evident.

These interpretations carry some methodological caveats. The chemical processing required for iterative expansion limits antibody compatibility, and because the antibodies validated for this protocol share the same host species, simultaneous co-localization of rhodopsin and peripherin-2 on individual discs was not feasible; the labeling patterns observed are nonetheless fully consistent with established biochemical evidence for each protein. The rhodopsin coverage estimate incorporates literature-derived geometric parameters and correlative electron microscopy was not performed on expanded specimens, but the close agreement between measured disc periodicity and cryo-electron tomography reference values confirms expansion isotropy at the relevant length scale, supporting the internal consistency of the quantitative measurements. Finally, disease-associated changes were assessed at a single early time point in the RCS model, chosen to capture the stage at which outer segment disruption is evident while sufficient structural integrity remains for single-disc quantification.

Collectively, these results establish iterative ultrastructure expansion microscopy as a platform for interrogating photoreceptor molecular organization at the single-disc level. This approach bridges a longstanding gap between the molecular specificity of fluorescence imaging and the structural resolution of electron microscopy, demonstrating that protein distributions within individual membrane compartments can be quantitatively resolved on conventional confocal hardware. The findings settle a persistent discrepancy in rhodopsin coverage estimates, provide direct molecular evidence for peripherin-2 within incisures, and uncover compartmentalized nanoscale pathology that precedes gross structural degeneration. Several concrete avenues extend from this work. Coupling iU-ExM with STED microscopy could push effective resolution below 5 nm, while multi-color labeling with antibodies raised in distinct host species would enable simultaneous mapping of multiple proteins within individual discs, directly addressing the co-localization constraint noted above. Extension to disease models harboring primary disc protein mutations and to longitudinal time points would test whether the compartmentalized vulnerability pattern generalizes beyond the RCS model. More broadly, the core principle demonstrated here, resolving molecular identity within structures too densely packed for conventional optics, is immediately transferable to ciliopathies affecting diverse cell types, synaptic nanodomains, and other tightly organized membrane systems where ultrastructural and molecular information have remained fundamentally uncoupled.

## Materials and Methods

### Experimental Design

#### Animal models and ethical approval

All experimental procedures were performed in accordance with standard European ethical guidelines (European Community Directive 2010/63/EU), approved by the local institutional committee, and conducted in compliance with the Association for Research in Vision and Ophthalmology (ARVO) Statement for the Use of Animals in Ophthalmic and Vision Research. Pigmented Long-Evans rats (wild-type) and Royal College of Surgeons (RCS-p+/Lav) rats were bred and housed under a 12-hour light/dark cycle with ad libitum access to food and water. Animals were euthanized at postnatal day 30 (P30) by CO₂ inhalation followed by cervical dislocation. Animals of both sexes were used; sex was not considered as a biological variable in this study, as sample sizes were optimized for methodological validation rather than sex-stratified comparisons.

#### Tissue Preparation

Eyes were enucleated and immediately punctured at the ora serrata using a 30-gauge needle to facilitate fixative penetration. Punctured eyes were immersed in 4% paraformaldehyde (PFA) in phosphate-buffered saline (PBS, pH 7.4) for 30 minutes at room temperature. Eyes were cryoprotected by immersion in 30% (w/v) sucrose in PBS overnight at 4°C and embedded in Optimal Cutting Temperature (OCT) compound (Tissue-Tek), snap-frozen, and cryosectioned at 10 µm thickness onto Superfrost Plus glass slides. Sections were stored at −20°C until processing for expansion microscopy.

#### Iterative Ultrastructure Expansion Microscopy (iU-ExM)

iU-ExM was performed following the protocol of Louvel et al. (*31*) with modifications for retinal cryosections. The procedure comprises two sequential rounds of hydrogel embedding and expansion, with intermediate immunolabeling between expansion steps. The complete protocol is described below.

1. Anchoring. Cryosections were incubated in anchoring solution (1.4% formaldehyde (FA), 2% acrylamide (AA) in 1× PBS) overnight at 37°C. Extended overnight incubation, rather than the standard 3 hours used for cultured cells, was employed to ensure thorough reagent penetration into the tissue.
2. First gelation. To enable homogeneous monomer diffusion throughout the tissue, sections were first incubated for 45 minutes in activated monomer solution (19% (w/v) sodium acrylate (SA), 10% (w/v) AA, 0.1% (w/v) N,N’-(1,2-dihydroxyethylene)bisacrylamide (DHEBA), 0.25% (v/v) TEMED and 0.25% (w/v) APS in 1× PBS). The DHEBA cleavable crosslinker, rather than the conventional non-cleavable N,N’-methylenebisacrylamide (BIS), is essential for the iterative approach as it permits selective dissolution of the first gel after re-embedding. A coverslip was placed on top to ensure uniform gel thickness. Polymerization proceeded for 45 minutes on ice followed by 1 hour at 37°C.
3. Denaturation. The gel was detached in denaturation buffer (200 mM SDS, 200 mM NaCl, 50 mM Tris, pH 6.8) under gentle agitation, then transferred to a 2 mL microcentrifuge tube with 1.5 mL of fresh buffer and incubated at 85°C for 2 hours. Temperature was monitored externally, as temperatures below 85°C cause anisotropic expansion while temperatures above 85°C degrade the DHEBA crosslinker. This denaturation regime, combined with the iterative expansion strategy, preserved outer segment disc membranes, which were not resolved in previous single-round expansion microscopy of photoreceptor tissue (*19, 32*).
4. First expansion. The gel was expanded in deionized water (ddH₂O), exchanging water every 20–30 minutes until expansion reached a plateau (∼5–6-fold linear expansion).
5. Intermediate immunolabeling. The first-expanded gel was shrunk in 1× PBS. Primary antibodies were diluted in 2% BSA in PBS at concentrations specified below and incubated overnight at 4°C, followed by three 15-minute washes with 0.1% Tween-20 in PBS (PBS-T). Secondary antibodies were incubated for 6 hours at 37°C in 2% BSA in PBS, followed by three 15-minute PBS-T washes. The gel was briefly re-expanded in ddH₂O to verify labeling quality.
6. Neutral gel embedding. Labeled gels were trimmed to ∼1 cm^2^ pieces and incubated 25 minutes on ice with activated neutral gel solution (10% AA, 0.05% DHEBA, 0.1% APS/TEMED in ddH₂O). The gel piece was placed on a glass slide, covered with a 22 × 22 mm coverslip, and polymerized for 1 hour at 37°C.
7. Re-anchoring. The neutral-gel-embedded specimen was incubated in anchoring solution (1.4% FA, 2% AA in PBS) overnight at 37°C under gentle agitation, followed by a 30- minute PBS wash. This step covalently tethers the antibody fluorescence signal to the second gel network and prevents signal loss during dissolution of the first gel.
8. Second gelation. The re-anchored gel was equilibrated by incubation on ice 25 minutes with second monomer solution (19% SA, 10% AA, 0.1% BIS, 0.1% TEMED/APS in PBS). The gel was covered with a coverslip and polymerized for 1 hour at 37°C. BIS (non-cleavable) crosslinker ensures this gel remains intact during subsequent dissolution.
9. Dissolution. The gel assembly was immersed in 200 mM NaOH for 1 hour at room temperature under agitation, selectively cleaving DHEBA crosslinks in the first and neutral gels. The gel was washed with PBS until pH returned to ∼7.0.
10. Final expansion. The second gel was expanded in ddH₂O, exchanging water until expansion plateaued, yielding a total linear expansion of approximately 20-fold.

Two features of the intermediate immunolabeling strategy are central to this study. First, at the first-expansion stage (∼5×), structures separated by 35 nm in native tissue are already spaced by ∼175 nm, enabling efficient antibody penetration into densely packed compartments such as photoreceptor discs. Post-expansion staining of the iteratively expanded second gel was not employed, as antibody diffusion into the dense gel network produces heterogeneous and weak labeling. Second, while the absolute linkage error introduced by primary/secondary antibody complexes remains constant (∼20–30 nm), the subsequent second expansion reduces the effective linkage error to ∼4–6 nm in native tissue dimensions (*31, 33*) a reduction critical for resolving protein localization within individual discs separated by only 30–35 nm.

#### NHS-Ester Protein Density Mapping

N-hydroxysuccinimide (NHS) ester-conjugated fluorophores react covalently with primary amine groups on proteins, generating a pan-proteomic density map analogous to electron microscopy contrast (*30, 31*). NHS-ester staining was performed on fully expanded second gels. Gels were briefly shrunk in 1× PBS and incubated with Alexa Fluor 594 NHS Ester (A20004, Thermo Fisher Scientific; 20 µg/mL in PBS) for 1.5 hours at room temperature under gentle agitation, followed by three washes of at least 30 minutes each with PBS. Gels were re-expanded in ddH₂O before imaging. When performed on gels that have undergone intermediate immunolabeling, the NHS-ester dye may additionally label primary and secondary antibodies, potentially contributing non-specific background at antibody-stained structures.

#### Primary and secondary antibodies

Primary antibodies: mouse monoclonal anti-rhodopsin (clone RET-P1, O4886, Sigma-Aldrich; 1:250), recognizing an intradiscal N-terminal epitope; mouse monoclonal anti-peripherin-2 (clone 6B10.1, MABN293, Sigma-Aldrich; 1:200); mouse monoclonal anti-α/β-tubulin (clone DM1A, T5168, Sigma-Aldrich; 1:250). Secondary antibodies: Aberrior STARGREEN and Aberrior STARRED goat anti-mouse (both 1:400). All dilutions were prepared in 2% BSA in PBS. Conventional primary/secondary antibody complexes were used throughout; smaller affinity reagents (nanobodies, Fab fragments) were not employed. Because all three primary antibodies derive from the same host species (mouse), simultaneous co-localization of rhodopsin and peripherin-2 on individual discs was not feasible. Although validation in peripherin-2-deficient tissue or with a second antibody clone recognizing a distinct epitope was not performed, several internal criteria support labeling specificity under iU-ExM conditions: peripherin-2 signal recapitulates the established hollow cylindrical rim pattern, is absent from the rhodopsin-occupied lamellar domain, and the 32 nm rhodopsin-free peripheral zone corresponds precisely to the peripherin-2-positive rim. This complementary exclusion pattern between two independently labeled proteins argues against off-target binding or denaturation-induced mislocalization.

#### Gel mounting and sample preparation for imaging

For all stages of expansion (1st or 2nd), gels were cut with a razor blade into squares to fit in 22 mm imaging chamber. The excess of water was carefully removed using absorbent tissue, care was taken to avoid gel desiccation. Next, the gel was placed on a Poly-D-Lysine coated 22 mm Mattek glass bottom dishes to prevent drifting.

#### Image Acquisition

Images were acquired using an Abberior Facility Line STED/confocal microscope equipped with 10x/0.4 NA air, 60×/1.4 NA oil-immersion and 60×/1.3 NA silicone-immersion objectives. Z-stacks were acquired at 0.25 µm (standard) or 0.1 µm (high-resolution) axial step sizes. Images were deconvolved using Huygens Professional software (Scientific Volume Imaging).

#### Expansion Factor Determination

For iteratively expanded gels, direct gel measurements are unreliable due to cumulative variations across expansion steps (*31*). Linear expansion factor was therefore determined for each experiment independently by comparing photoreceptor nuclear cross-sectional area between non-expanded and expanded samples, following the biological ruler approach established for tissue-based iU-ExM. Photoreceptor nuclei were selected as the reference structure due to their low dimensional variability across the retinal section, enabling precise estimation. Expansion factors varied across experimental sessions (WT: 19.6 ± 2.8×, n = 158 nuclei; RCS: 21.2 ± 1.8×, n = 93 nuclei; pooled mean: 20.2 ± 2.65×), reflecting differences in reagent freshness between sessions rather than genotype-dependent tissue properties. Because each experiment’s individually determined expansion factor was applied to correct all measurements within that dataset to native tissue dimensions, batch-to-batch variation does not propagate into corrected values. Internal validation confirmed expansion accuracy by two independent criteria: (i) autocorrelation-derived inter-disc periodicity (35 ± 2.8 nm) matching established cryo-electron tomography values (*1, 3*), confirming nanometer-scale accuracy within the structures under investigation; and (ii) preservation of ninefold radial symmetry in centriolar appendages (Fig. 4C-D), indicating isotropic expansion across compartments. The concordance between nucleus-derived expansion factors (operating at micrometer scale) and disc periodicity measurements (operating at nanometer scale) provides convergent validation across two orders of magnitude in spatial dimension.

#### Image Analysis

Inter-disc spacing. Measured from intensity profiles perpendicular to the outer segment axis. Direct peak-to-peak distances were obtained from individual profiles; one-dimensional autocorrelation analysis with angular correction (±10°) provided robust periodicity estimates.

Rhodopsin radial coverage. Quantified from radial intensity profiles measuring rhodopsin-occupied diameter relative to total outer segment diameter (plasma membrane to plasma membrane). The peripheral exclusion zone was computed by subtracting literature-derived plasma membrane-to-disc edge spacing (∼25 nm) and membrane thickness (∼7 nm) from the measured plasma membrane-to-rhodopsin signal distance (2). Coverage was calculated as the ratio of rhodopsin-occupied to total lamellar domain, assuming isotropic radial expansion (Fig. S1D). The rhodopsin coverage estimate incorporates literature-derived parameters for plasma membrane-to-disc edge spacing, introducing systematic uncertainty not captured in the reported variability.

Disc morphometry. Measured from peripherin-2-labeled en face views using particle analysis in ImageJ/FIJI for area, perimeter, and roundness (4 × area / π × major axis²). Incisure depth was measured manually as the inward extent of peripherin-2 signal from the disc rim toward the disc center. The incisure membrane perimeter was calculated as the difference between total disc perimeter and perimeter excluding incisures. This is because the membrane lines both arms of the incisure fold, this value is approximately twice the incisure depth. Measurements were restricted to clearly resolved structures exhibiting characteristic morphology. Correlative electron microscopy was not performed on expanded specimens; incisure assignments therefore rely on fluorescence-based geometric criteria.

Connecting cilium morphometry. Quantified from serial cross-sections at 12.5 nm effective axial intervals (0.25 µm acquisition step corrected for expansion factor). Area, perimeter, and roundness (4 × area / π × major axis²) were measured at each position. Transition zone onset was identified using derivative analysis of the smoothed area-versus-position curve. The first derivative (slope) was insufficient for onset detection because the gradual transition from near-zero baseline values made identification of the departure point threshold-dependent. The second derivative (curvature) identified the principal inflection point, position of maximum acceleration, but not the onset of departure from linearity. Third-derivative analysis resolved this by detecting the point at which curvature itself begins to increase: the first significant peak in the third derivative marks the earliest departure from the baseline linear regime and was used to define transformation zone onset. The principal inflection point was independently identified from the second-derivative maximum. Curves were smoothed prior to differentiation to suppress high-frequency noise amplification inherent to successive differentiation. The same smoothing parameters and derivative thresholds were applied identically to wild-type and RCS datasets. Axial positions were defined relative to the outer segment boundary, identified as the first contiguous rhodopsin-positive disc band.

Centriolar appendages. DAP and sDAP structures were identified from NHS-ester-labeled en face views displaying ninefold radial symmetry. Diameters, inter-appendage spacing, and positions relative to the basal body were measured from en face and longitudinal views.

RCS sample selection. Quantitative analysis was restricted to outer segments retaining sufficient structural integrity for autocorrelation-based disc resolution. Severely disrupted outer segments were excluded, limiting conclusions to early-to-moderate pathology in the analyzable subpopulation. Per-experiment expansion factor correction was applied independently to each dataset. Several internal criteria argue against differential expansion artifacts in RCS tissue: disc dimensions (area and perimeter) were equivalent between genotypes, and centriolar appendage architecture in the inner segment showed no significant differences. Correlative electron microscopy was not performed on expanded specimens; while preserved disc dimensions between genotypes and matching wild-type periodicity with cryo-electron tomography values provide internal validation, direct structural correspondence in diseased tissue remains to be established.

### Statistical Analysis

Data are presented as mean ± s.d. Comparisons between wild-type and RCS groups used unpaired two-tailed Student’s t-tests (normally distributed data) or Mann-Whitney U tests (non-normal distributions, assessed by Shapiro-Wilk test). Significance was defined as P < 0.05. Sample sizes represent measurements from at least three independent animals per genotype. All statistical analyses were performed using Python (SciPy, Matplotlib).

## Acknowledgments

We extend our gratitude to Esther Aguilar (FCRB-IDIBAPS) for their technical support.

## Author contributions

- Conceptualization: SM, PLA
- Methodology: SM
- Investigation: SM, EPP, JP
- Formal analysis: SM
- Visualization: SM
- Writing—original draft: SM
- Writing—review & editing: SM, JP, EPP, PLA
- Supervision: PLA
- Funding acquisition: PLA

## Data and materials availability

All data are available in the main text or the supplementary materials. The data supporting the findings of this study are available from the corresponding author. Analysis code is available at https://github.com/simone-mortal/Disc_Photo_iUExM_2026.

